# Effect of mounding, drainage and fertilization on CH_4_ fluxes and methane-cycling functional genes in waterlogged forest stands

**DOI:** 10.1101/339978

**Authors:** David J. Levy-Booth, Cindy E. Prescott, Susan J. Grayston

## Abstract

Site preparation techniques including mounding, drainage and nitrogen (N) fertilization can enhance seedling survival and site productivity, particularly in waterlogged, low-productivity forest stands. However, practices that alter soil conditions and site hydrology can lead to the unintended alteration of biogeochemical process rates, such as CH_4_ fluxes. This study sought to measure CH_4_ fluxes measured using static closed chambers at a sub-boreal spruce site and a coastal cedar-hemlock site that underwent mounding and drainage, respectively, to manipulate water table depth relative to planted seedlings, as well as fertilization. The abundance of methyl coenzyme M reductase (*mcrA*) gene found in methanogenic archaea and the particulate methane monooxygenase (*pmoA*) gene found in methane-oxidizing bacteria (MOB) were examined. The use of sulphate as a potential method to stimulate sulphate-reducing bacteria (SRB) and reduce methanogen activity was also investigated using the dissimilatory sulfite reductase β-subunit (*dsrB*) gene. qPCR was used to link *mcrA,pmoA* and *dsrB* gene abundance to soil factors and GHG fluxes. Mounding created hot-spots of CH_4_ emissions at the spruce site. Drainage improved soil aeration at the coastal cedar-hemlock site and reduced CH_4_ emission rates. Fertilization did not affect CH_4_ emissions from either site. CH_4_ rates were influenced by soil water content and *mcrA* abundance. Measurements of microbial functional genes can elucidate the effects of site preparation on GHG fluxes in waterlogged forest stands.

## Introduction

Elevated atmospheric concentrations of greenhouse gases (GHGs) are of major concern worldwide. Methane (CH_4_) is the second most-important GHG in terms of radiative climate forcing, its concentrations has increased by about 150% in the last two centuries to about 1.8 ppm (Forster et al., 2007). CH_4_ fluxes in forest and wetland soil is a major component of the global carbon (C) cycle. Boreal (1372 Mha) and temperate forests (1038 Mha) contain 272 and 119 Pg C in total, and are sequestering an additional 0.5 and 0.72 Pg C yr^−1^ respectively, of which 65% and 49% is stored in soil (Pan et al., 2011). Alterations to the soil C cycle can lead to the release of substantial stored carbon, which is exacerbated in wet forests due to the incomplete degradation of plant biomass carbon in anaerobic zones.

Forest site preparation such as drainage, mounding as well as nitrogen (N) fertilization are used to enhance seedling establishment and growth in wet forest ecosystems, and can increase site C sequestration through the accumulation of aboveground biomass and soil C (Laiho and Finér, 1996; Laiho and Laine, 1997; Canary et al., 2000; Johnson and Curtis, 2001; Bond-Lamberty et al., 2002; Blaško et al., 2013). However, site C sequestration due to site preparation and fertilization could be offset by increasing GHG fluxes through alterations to the microbial community (Jandl et al., 2007; Mojeremane et al., 2012). This study examines the microbial communities involved in CH_4_ fluxes in wet forest soil ecosystems.

Methanogens from phylum *Euyarchaeota* produce nearly all biogenic CH_4_ using a variety of metabolic pathways, though methanogenesis in terrestrial ecosystems is primarily hydrogenotrophic (CO_2_ + H_2_) or aceticlastic (acetate) (Conrad, 1999, 2005). Molecular characterization of methanogens target the *mcrA* gene, which encodes the methyl coenzyme M reductase enzyme common to all known methanogensis pathways (Luton et al., 2002). Aceticlastic methanogensis is responsible for about two-thirds of methane production in soil (Le Mer and Roger, 2001). Studies of methanogen community structure, including *mcrA* and 16S rRNA characterization, suggest that both hydrogenotrophic methanogens and aceticlastic methanogens are common in waterlogged forest soil (Kemnitz et al., 2004; Conrad, 2005; Frey et al., 2011; Kanokratana et al., 2011). Methanogen diversity (measured by *mcrA* and 16S rRNA) and methanogenesis in forest soil are weakly but positively influenced by soil temperature (Fey and Conrad, 2000; Krause et al., 2013), and strongly and positively correlated with soil water content (Ullah et al., 2009; Hartmann et al., 2014). Management practices that change these soil parameters can greatly alter CH_4_ flux from forest soil (Fey and Conrad, 2000; Watanabe et al., 2009; Ma et al., 2011; Angel et al., 2012; Hartmann et al., 2014). Methanogensis has very low energy yields (ΔG^°^’= −131 and −136 kJ for hydrogenotrophic and aceticlastic pathways, respectively) and generally occurs in very low redox potential soils, as methanogens are generally out-competed for acetate and protons by other biological reducers, e.g., sulphate-reducing bacteria (SRB) (ΔG^°^’= −152.2 kJ) (Thauer et al., 1989; Muyzer and Stams, 2008). It is, therefore, predicted that SO_4_-S fertilization of waterlogged soil can stimulate the SRB (characterized using the dissimilatory sulfite reductase β-subunit *(dsrB)* gene) and suppress aceticlastic CH_4_ production.

Methane-oxidizing bacteria (MOB) in temperate upland forest soil soil provide a net sink of atmospheric CH_4_ (Adamsen and King, 1993; Dutaur and Verchot, 2007; MacDonald et al., 1996; Krause et al., 2013), and CH_4_-uptake in soils can account for between 15 and 45 Tg CH_4_ uptake yr^−1^ (Wuebbles and Hayhoe, 2002). The MOB contain either soluble or particulate methane monooxygenase (PMO) enzymes to oxidize CH_4_, which are encoded by the *mmoX* and *pmoA* genes, respectively. Nearly all MOB (except genera *Methyloferula* and *Methylocella)* contain *pmoA,* the structure and abundance of which is influenced negatively by soil water content and weakly but positively by soil temperature, pH and forest type (Dunfield, 2007; Kolb, 2009; Shrestha et al., 2012). The use of molecular markers for methanogens and MOB can elucidate effects of site preparation and management on the organisms driving CH_4_ fluxes in forest ecosystems.

Mechanical site preparation (i.e., ditch drainage and excavator mounding) can alter site hydrology, soil moisture and temperature, creating planting sites ideal for economically-important tree species. Drainage leaves soil structure and stand vegetation relatively undisturbed, while mounding can disrupt soil structure, bury forest floor layers and remove competing vegetation (Åkerström and Hånell, 1996; Örlander et al., 1998). Alterations to the soil environment following site preparation can enhance litter decomposition, increase N mineralization and nitrification, increase soil respiration and reduce CH_4_ emissions (Martikainen et al., 1995; Lundmark-Thelin and Johansson, 1997; Smolander et al., 2000; Minkkinen et al., 2002; von Arnold et al., 2005; Mojeremane et al., 2012).

Fertilization can increase soil organic C and N concentrations (Smolander et al., 2000; Johnson and Curtis, 2011; von Arnold et al., 2005; Jandl et al., 2007), and can increase (Hasselquist et al., 2012; Mojeremane et al., 2012) or decrease (Liu and Greaver, 2009; Janssens et al., 2010; Krause et al., 2013) respiration from forest soil. Decreases in CH_4_ uptake in upland soil or increases in CH_4_ emissions from wetland soil following N deposition or fertilization can result from NH_4_^+^ saturating the binding sites for CH_4_ in PMO, suppressing CH_4_ oxidation (Basiliko et al., 2009; Jassel et al., 2011), possibly due to evolutionary similarities between PMO and ammonium monooxygenase enzymes from nitrifying bacteria (Bédard and Knowles, 1989; Holmes et al., 1995; Purkhold et al., 2000; Bodelier and Laanbroek, 2004). However, N fertilization has also been shown to increase CH_4_ flux rates as a result of decreased CH_4_ uptake (Castro et al., 1994; Liu and Greaver, 2009; Mojeremane et al., 2012), suggesting that the role of N addition to forest soils in regulating CH_4_ flux is not yet fully understood (Gundersen et al., 2012). Characterization of the soil environment and microbial community in a variety of forest ecosystems can identify the underlying causes of CH_4_ flux differences following site preparation and N fertilization.

This study seeks to use quantitative PCR (qPCR) of the *mcrA, pmoA* and *dsrB* genes to estimate the response of the methanogen, MOB and SRB communities, respectively, in two regenerating waterlogged forest stands subject to mounding, drainage and fertilization. We attempt to link these functional groups to CH_4_ fluxes measured using static closed chambers to better understand the importance of the microbial community in determining GHG fluxes from managed forest stands.

## Materials and Methods

### Field sites

The effects of fertilization and site preparation (mounding and drainage) on GHG flux, and on the abundance of methanogenic archaea, MOB and SRB was investigated at two waterlogged sites in British Columbia (B.C.), Canada. Field site descriptions are provided in Levy-Booth et al. (2016). Briefly, mounding treatments were installed at Aleza Lake Research Forest (ALRF), in the wk1 variant of the sub-boreal spruce (SBS) biogeoclimatic zone (Green and Klinka 1994), near Prince George, B.C. Soils at ALRF are Orthic Gleyed Luvisols, Orthic Luvic Gleysols and Ortho Humo-Ferric Podzols. The 70-year old second-growth stand of interior hybrid spruce *(Picea engelmannii* x *glauca)* and subalpine fir *(Abies lasiocarpa)* were harvested in Feburary 2011 and replanted with interior hybrid spruce on June 6, 2012. Excavator mounding took place in August 22, 2011. Fertilizer was applied at a final formulation of 200 kg N, 100 kg P, 100 kg K, and 50 kg S ha^−1^ on June 26, 2012.Treatment plots were organized in a complete-block design, with two blocks containing each of the four treatments (unmounded/unfertilized, unmounded/fertilized, mounded/unfertilized, mounded/fertilized). In mounded plots, the tops of mounds as well as the hollows were sampled. Sampling took place on June 23, 2011, June 28, July 17, August 24, October 18, 2012 and June 13, 2013, corresponding to pre-mounding and 24 hours, 1 month, 2 months, 4 months and 1 year following fertilization. The Suquash Drainage Trial (SDT) is located near the Salal Cedar Hemlock Integrated Research Program (SCHIRP) research site installed by Western Forest Products Inc. between the towns of Port Hardy and Port McNeill on northern Vancouver Island, B.C. The SDT site is located in the vm1 subzone of the Submontane Very Wet Maritime Coastal Western Hemlock ecozone (CWHvm1). Soil is Humo-Ferric Podzols with mor humus. The original 22 ha western redcedar *(Thuja plicata*) and shore pine *(Pinus contorta* var. *contorta*) stand underwent harvesting and slash-burning in 1993 and 1994, respectively, and planted with western redcedar *(Thuja plicata)* in 1995. Three 120 m x 45 m treatment plots containing 5 drainage ditches from a 1997 installation were used in this study. Operational fertilization of 225 kg N and 75 kg P ha^−1^ was conducted in 2006. Undrained control plots were selected at least 60 m away from each ditched area to avoid the effects of ditching on subsurface drainage, which extended 15 m from each drainage ditch (van Niejenhuis and Barker, 2002). One of two 30 x 10 m transects in each drained or undrained plot was on July 25, 2012 at the same formulation used in ALRF. Plots were organized in complete-block design and included the following treatments: drained/fertilized, drained/fertilized, drained/unfertilized and drained/unfertilized. Soil sampling at SDT for microbial gene analysis took place on July 27, 2012 (Jul-12), August 29, 2012 (Aug-12), October 25, 2012 (Oct-12), July 3, 2013 (Jul-13) and September 12, 2013 (Sep-13). GHG measurements did not occur in Oct-12 and soil chemical factors were not measured in Sept-13.

### Soil sampling and preparation

At ALRF and SDT mineral soil (Ae and some Bf horizons; ~ 0-5 cm) were sampled from three and two randomly chosen locations in each treatment plot, respectively. Volumetric soil moisture was measured using a TH_2_O^TM^ portable moisture probe (Dynamax Inc., Houston, U.S.A.) and gravitational soil moisture was measured by oven drying field moist soil. Results of Total C, Total N, NO_3_-N, NH_4_-N and pH analysis are described in Levy-Booth et al. (2016). In this study, 10 g dry soil was analyzed for total S and SO_4_-S by the British Columbia Ministry of Forests, Lands and Natural Resources Operation Analytical Laboratory (Victoria).

### Field measurement and gas chromatography analysis of CH_4_ fluxes

Measurement of *in situ* GHG flux at ALRF and SDT took place as described in Basiliko et al. (2009) and in Levy-Booth et al. (2016). Briefly, three closed PVC chambers were installed on collars buried about 5 cm in the soil in each of two treatment plots at ALRF, and two chambers were installed in each of three treatment plots at SDT. Six ml of chamber headspace were removed and inserted into pre-evacuated 5 ml Exetainers^®^ (Labco Ltd., Lampeter, UK) every 15 minutes for one hour. Gas samples were measured on an Agilent 5890 series II gas chromatograph (Agilent Technologies, Santa Clara, U.S.A.) equipped with a flame ionisation detector (FID) set at 300^°^C. The FID carrier gas was helium with a flow rate of 14 ml min^−1^. Standards for gas chromatography used 4, 2, 1 and 0.67 ppm CH_4_.

### Nucleic acid extraction and quantitative PCR

DNA was extracted from 0.25 g dry soil using the MoBio PowerClean soil DNA isolation kit and quantified using the Quant-iT^TM^ PicoGreen^®^ dsDNA assay (Life Technologies Corp., Carlsbad, U.S.A.). qPCR used Power SYBR^®^ Green PCR Master Mix (Life Technologies Corp., Carlsbad, U.S.A.) in the Applied Biosystems^®^ StepOnePlus^TM^ real-time PCR system. Gene copy numbers were expressed as copy number g^−1^ soil (dry weight (dw)). *McrA* and *pmoA* quantification are described in Christiansen et al. (2016) using primers from Luton et al. (2002) and Bourne et al. (2001), respectively, and amplification protocols in Frietag et al. (2010). Standard curves for calibration of *mcrA* qPCR were created using triplicate 10-fold dilutions from 10^3^ to 10^8^ copies of clonal 420-bp *mcrA* segments from *Methanolinea mesophila,* amplified from soil near the SDT site as in Christiansen et al. (2016). Standard curves for *pmoA* were created using triplicate 10-fold dilutions from 10^3^ to 10^8^ copies of *Methylococcus capsulatus* genomic DNA. *DsrB* qPCR used forward (Dsr2060f, 5’-CAACATCGTYCAYACCCAGGG-3’) and reverse (Dsr4r, 5’-GTGTAGCAGTTACCGCA-3’) primers from Geets et al. (2006) at concentrations of 0.5 μM each. QPCR included an initial denaturation of 5 min at 95^°^C and 40 cycles of 95^°^C denaturation for 30 s, 55^°^C annealing for 45 s, 72^°^C extension for 45 s, after which fluorescence was measured. Standard curves for *dsrB* were created using triplicate 10-fold dilutions from 10^2^ to 10^8^ copies of *Desulfosporosinus orientis* and *Desulfomicrobium baculatum* genomic DNA.

### Statistical analysis

Statistical analysis was performed using in R v. 2.15.3 (R Core Team, 2013). Data were fitted with the linear mixed-effects model and subject to two-factor analysis of variance (ANOVA) using the *lme* and *anova* functions in the *nlme* and car packages, respectively, with fertilization and mounding/drainage as fixed effects and blocking as a random effect. qPCR data were analyzed as log10 values and plotted using 25%-75% quartile boxplots with SigmaPlot 11.0 (Systat Software, Inc., San Jose, CA). GHG data that violated ANOVA assumptions were logarithmically transformed and qPCR data that violated assumptions were analyzed as normal values. Unconstrained, exploratory ordination was carried out using principal component analysis (PCA) on scaled parameters with the *prcomp* function for ordination by singular value decomposition and visualized using the *ggbiplot* package in R. Forward selection of significant soil parameters was performed using permutational testing with 1000 permutations. Principal coordinate of a neighbour matrix (PCNM) analysis (Borcard and Legendre, 2002; Borcard et al., 2004) was used to generate variables for spatial structure using the *PCNM* function in the *PCNM* package in R. The geographic locations of sample sites were transformed to Cartesian coordinates using *geoXY* function in the *SoDA* package in R prior to PCNM. PCNM variables were tested using Moran’s I statistic to select significant and positive variables, which were then forward selected against spatially detrended dependant variables or matrices using permutational testing with 1000 permutations of the reduced model, ensuring that R^2^-adjusted of the forward-selected models did not exceed the R^2^-adjusted of the non-selected models. Canonical variation partitioning using RDA (Borcard et al., 1992; Ramette and Tiedje, 2007; Bru et al., 2011) was carried out to allocate dependant variable or matrix variance uniquely explained by each soil parameter, or groupings of parameters, by constraining linear partial regression by all other variables. Bonferroni-corrected p-values were used for multiple comparisons.

## Results

### Site preparation altered forest soil CH_4_ flux rates

Site preparation had a significant but inconsistent effect on CH_4_ flux rates in both ALRF and SDT sites. At ALRF, the highest rates CH_4_ efflux were measured following fertilization on Jul-12 (Figure 1a). At this date, CH_4_ efflux in mound hollows rates was 422.4 ± 176.6 and 1171.3 ± 368.8 μg CH_4_ m^−2^ h^−1^ in unfertilized and fertilized plots, respectively. By Aug-12—the warmest, driest date—79% of chambers demonstrated CH_4_ uptake. Mounding appeared to reduce CH_4_ flux rates after one year, owing primarily to the reduction of efflux in exposed mounds. There was consistent CH_4_ uptake measured in the drained plots at SDT, particularly in chambers located in unfertilized plots (Figure 1b). While there was CH_4_ uptake in all plots throughout the course of the field measurements, the undrained control plots had significantly greater emissions throughout the experiment (Jul-12, Aug-12 and Jun-13). SDT fluxes did not suggest seasonal or fertilization effects, having less temporal fluctuations than at ARLF.

**Figure 1.**
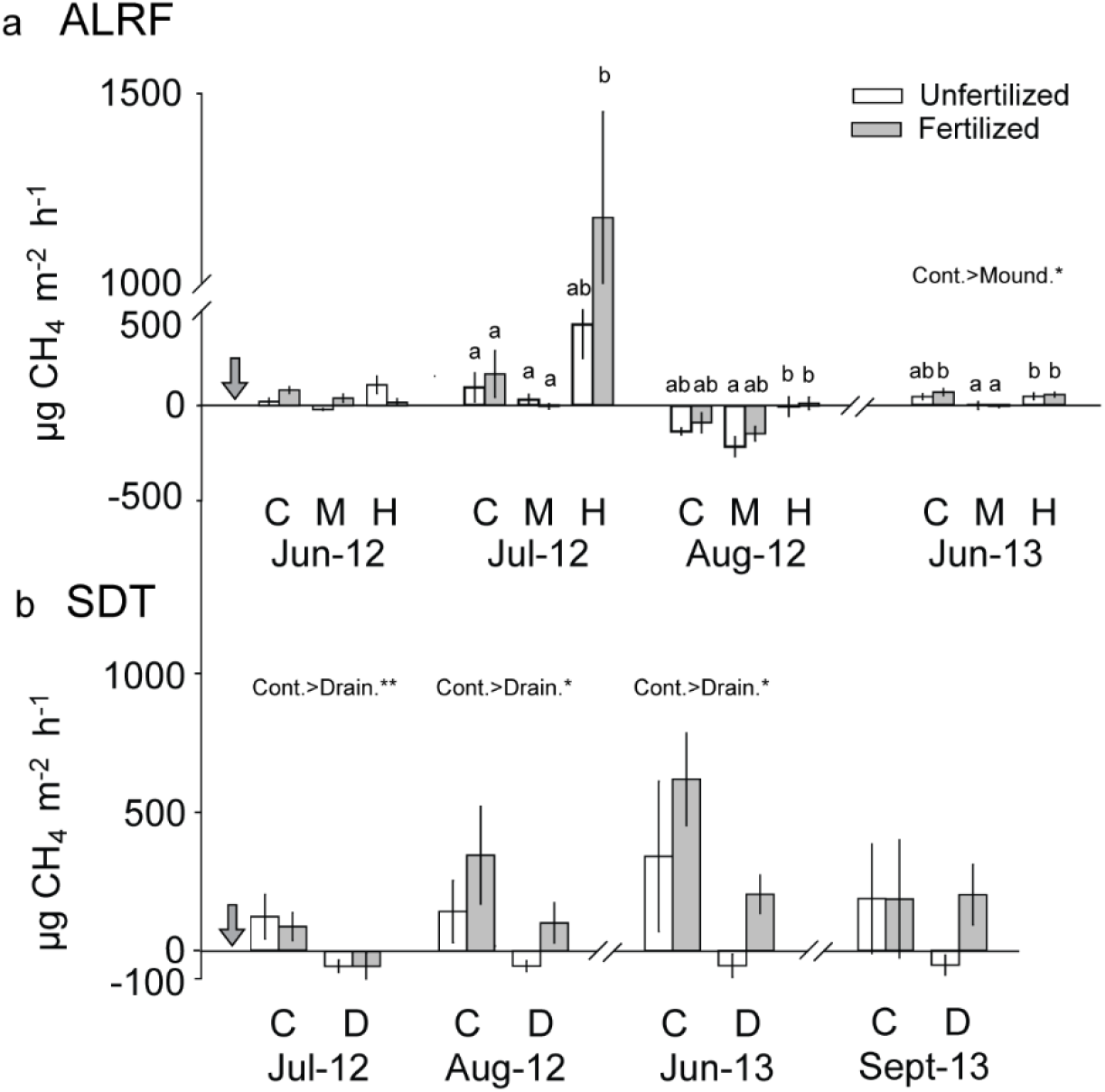
a) CH_4_ fluxes emissions from a) undisturbed control (C) and mounded plots (M, mounds; H, hollows) subject to fertilization at ARLF and b) undrained control (C) and drained (D) subject to fertilization at SDT. Shaded arrow shows time of fertilization. Error bars: SEM. N = 6. Treatment locations identified by different letters were significantly different at p < 0.05 following one-way ANOVA. Treatment effects and interactions following two-way ANOVA are provided if significant (*, p<0.05, **, p<0.01, ***, p<0.001).

### Fertilization transiently increased SO_4_–S concentrations

An important component of this study was to determine how fertilization with S affected the SRB and the methane-cycling community. Trace concentrations of total S were only measured at ALRF in the forest floor F and H layers (data not shown). However, SO_4_ was measured in almost all of the mineral samples taken from this location (Figure 2a). Treatment effects were measured at ARLF throughout the study, with a significant increase after 24 hours of fertilization. This effect was only observed in the mounded plots, which lacked an organic layer. As a result there was a significant interaction between mounding and fertilization at this date. The fertilization effect continued in Jul-12, although the SO_4_-S concentrations in the fertilization plots had decreased. SDT had trace amounts of total S in 52% of forest floor samples (data not shown) from both fertilized and unfertilized plots. In contrast to ALRF there were no treatment or locational effects of mounding and fertilization on SO_4_-S at SDT, with the exception of Jul-13, where fertilized drained soil samples had significantly greater SO_4_-S than control and drained unfertilized samples (Figure 2b). Soil SO_4_-S was greater at SDT than at ARLF, with concentrations reaching a peak of 90.1 ± 35.2 and 16.5 ± 4.2 μg g soil at these sites, respectively.

**Figure 2.**
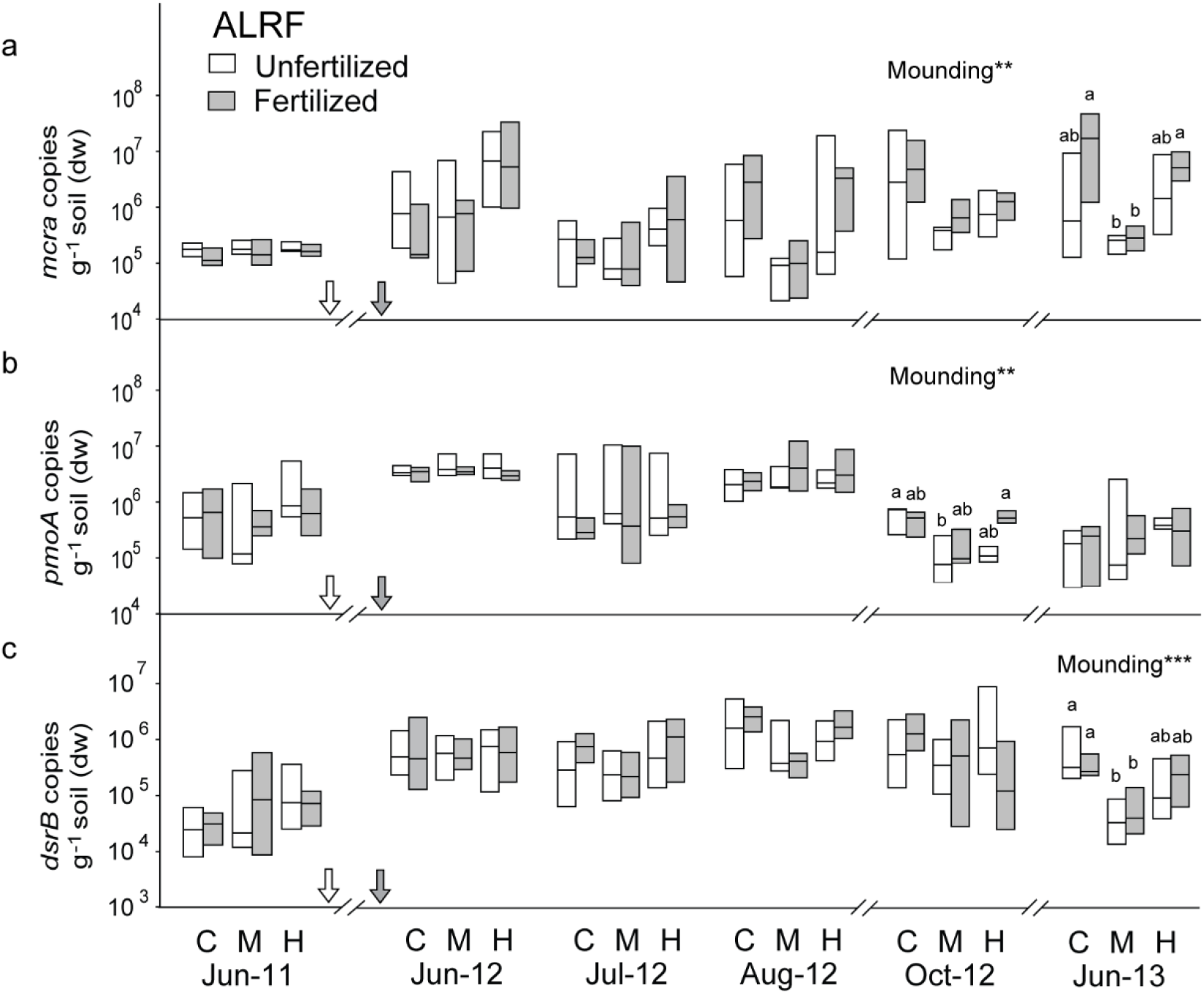
SO_4_-S concentration (mg kg^−1^) in soil from a) from undisturbed control (C) and mounded plots (M, mounds; H, hollows) subject to fertilization at ARLF, and b) control (C) and drained plots (D) subject to fertilization at SDT. Shaded arrow shows time of fertilization. ± SEM, n = 6. Treatment locations identified by different letters were significantly different at p = 0.05 following one-way ANOVA. Treatment effects and interactions following two-way ANOVA are provided if significant (*, p<0.05, **, p<0.01, ***, p<0.001).

### Microbial gene abundance altered by site preparation

*mcrA* abundance was affected by mounding, with significantly lower abundance in mounded plots in Oct-12 (Figure 3a). In Jun-13, as the mounded plots once again became waterlogged, fertilized mound hollows and unmounded plots had significantly greater *mcrA* abundance than mound tops. A consistent drainage effect on *mcrA* abundance was observed at SDT (Figure 4b). Undrained control plots had significantly greater *mcrA* abundance than drained plots in Aug-12 and Jul-13 and Sep-13. A significant effect of fertilization also occurred in Jul-13, one year following the treatment.

**Figure 3.**
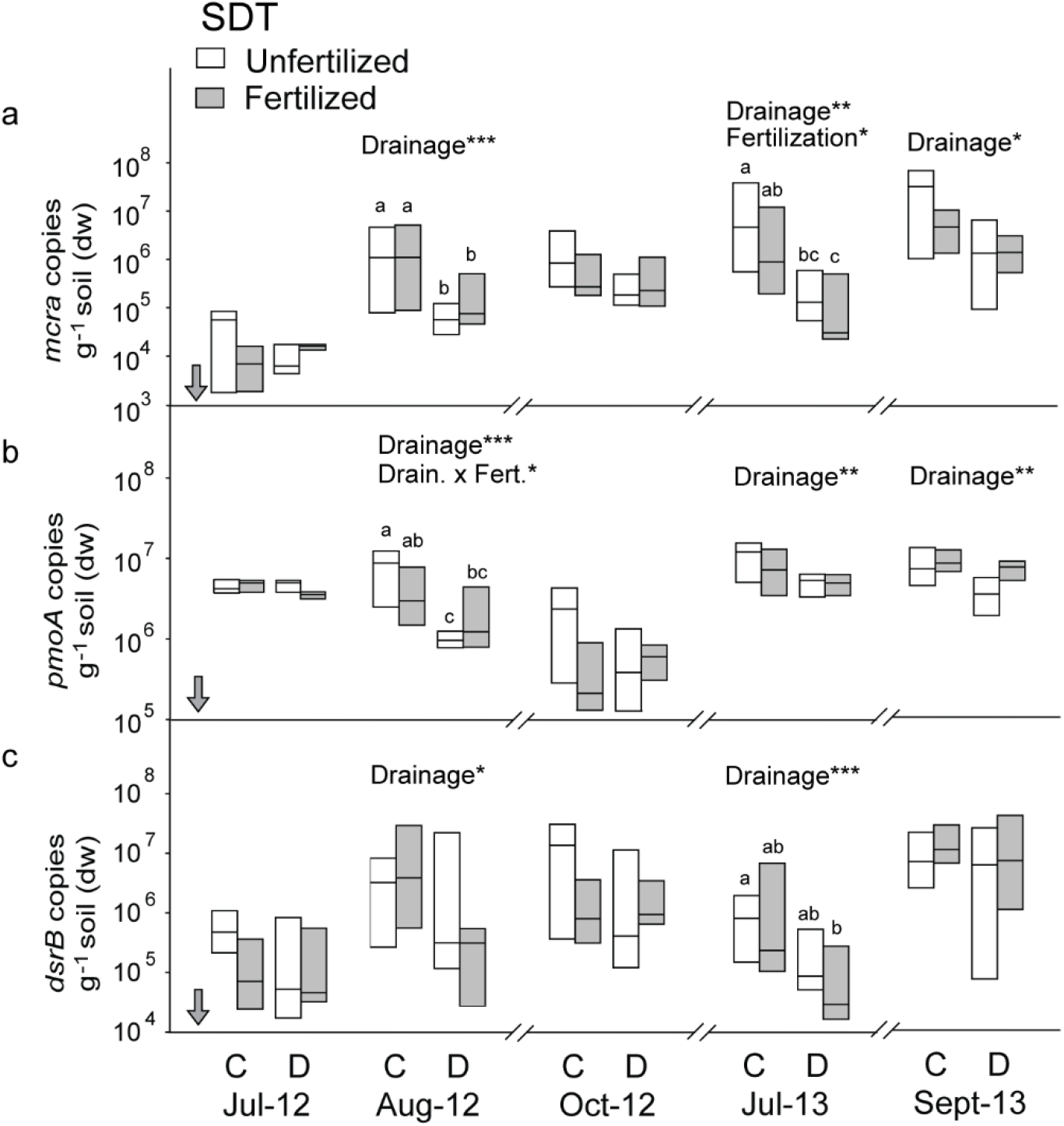
Abundance of a) *mcrA* genes, b) *pmoA* genes and c) *dsrB* genes in soil from undisturbed control (C) and mounded plots (M, mounds; H, hollows) subject to fertilization at ARLF. White arrow shows time of mounding, shaded arrow shows time of fertilization. Boxplots show median, 25% quartile and 75% quartile; n = 6. Treatment locations identified by different letters were significantly different at p < 0.05 following one-way ANOVA. Treatment effects and interactions following two-way ANOVA are provided if significant (*, p<0.05, **, p<0.01, ***, p<0.001).

**Figure 4.**
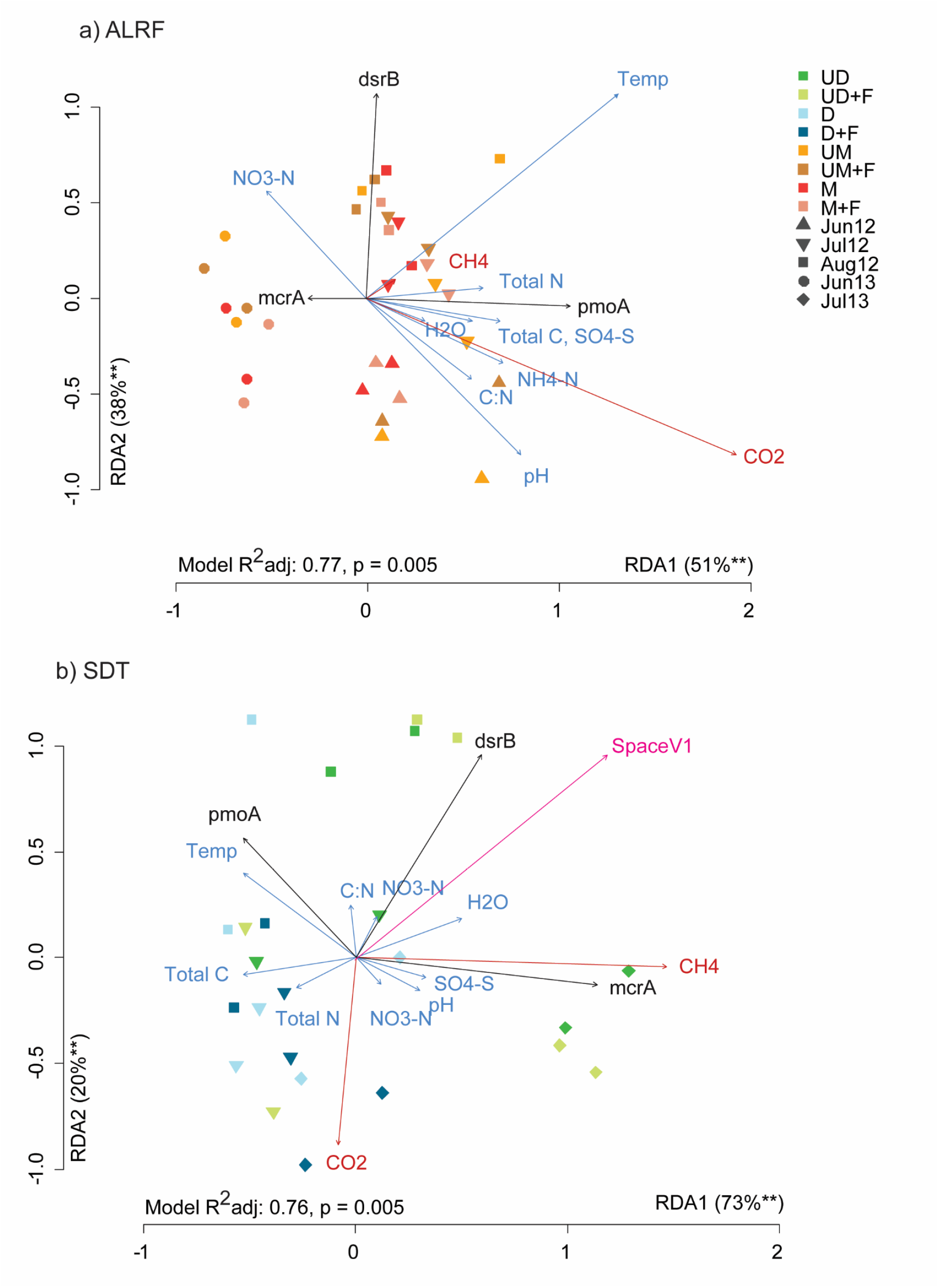
Abundance of a) *mcrA* genes, b) *pmoA* genes and c) *dsrB* genes in soil from undrained control (C) and drained (D) subject to fertilization at SDT. Shaded arrow shows time of fertilization. Boxplots show median, 25% quartile and 75% quartile; n = 6. Treatment locations identified by different letters were significantly different at p = 0.05 following one-way ANOVA. Treatment effects and interactions following two-way ANOVA are provided if significant (*, p<0.05, **, p<0.01, ***, p<0.001).

The abundance of *pmoA* was significantly greater in mounded plots than in unmounded plots in Oct-12 (Figure 3b). *PmoA* abundance in ALRF and SDT soil appeared to follow a seasonal trend. The lowest measured abundance (5.3 ± 0.2 *pmoA* copies g^−1^ soil (dw)) was measured in undrained SDT fertilized plots in Oct-12, which had a soil temperature of 8.1^°^C. Mirroring *mcrA* abundance, control plots had significantly greater *pmoA* abundance than drained plots in Aug-12 as well as Jul-13 and Sep-13. There was significant interaction between drainage and fertilization in Aug-12, as fertilization counteracted the effect of drainage on *pmoA* abundance.

*DsrB* abundance at the ALRF site was significantly lower in mounded plots in Jun-13 (Figure 3c), mirroring to *mcrA* abundance at this date. At SDT, *dsrB* was significantly lower in drained soil in Aug-12 and Jul-13. There was no increase of *dsrB* genes related to application of SO_4_-S fertilizer, nor were interactions between fertilization and drainage observed.

### Influence of site preparation and fertilization on soil parameters, gene abundance and GHG flux

Redundancy analysis (RDA) was used to determine the relationships between soil parameters and microbial gene abundance. In addition to the variables reported in this study, soil chemical variables, bacterial 16S rRNA gene abundance and CO_2_ flux rates from Levy-Booth et al. (2016) were included in this analysis. Forward-selected soil variables explained 77% of variation in ALRF gene abundances (Figure 5a). *McrA* abundance was negatively correlated with *pmoA* abundance, CO_2_ fluxes and soil temperature, but positively correlated with CH_4_ flux rates (Supplementary Table 1). *PmoA* abundance was strongly and positively correlated with CO_2_ fluxes and soil temperature. Canonical variation partitioning using partial regression was used to test variables that explain variation in gene abundances and GHG fluxes. Note that the sum of variation explained by individual variables will not equal the total model variation, as additive or subtractive effects can occur. Bacterial 16S rRNA gene abundance was explained most strongly by total soil carbon and nitrate concentration. Soil temperature explained the most variation in *McrA, pmoA* and *dsrB* abundance, with soil water content and NO3-N concentrations explaining additional variation. *PmoA* was also significantly explained by *mcrA* abundance. In turn, CO_2_ fluxes were significantly explained only by soil temperature and NO3-N concentrations, while *mcrA* abundance, bacterial 16S rRNA gene abundance and NO3-N explained CH_4_ fluxes, albeit only 8.6% of variation in CH_4_ fluxes could be explained at ALRF. However, these data indicate that the abundance of microorganisms containing *mcrA, pmoA* and *dsrB* genes can fluctuate with temperature, and are influenced by nitrate availability. Further, CH_4_ fluxes, while poorly explained by soil variables were most-strongly linked to the population size of *mcrA* containing organisms at ALRF. While fertilization with NO3-N and SO_4_-S was thought to suppress methanogenesis in forest soil, fertilization may lead to a stimulation of these populations through the alleviation of nitrogen limitation.

**Table 1.**
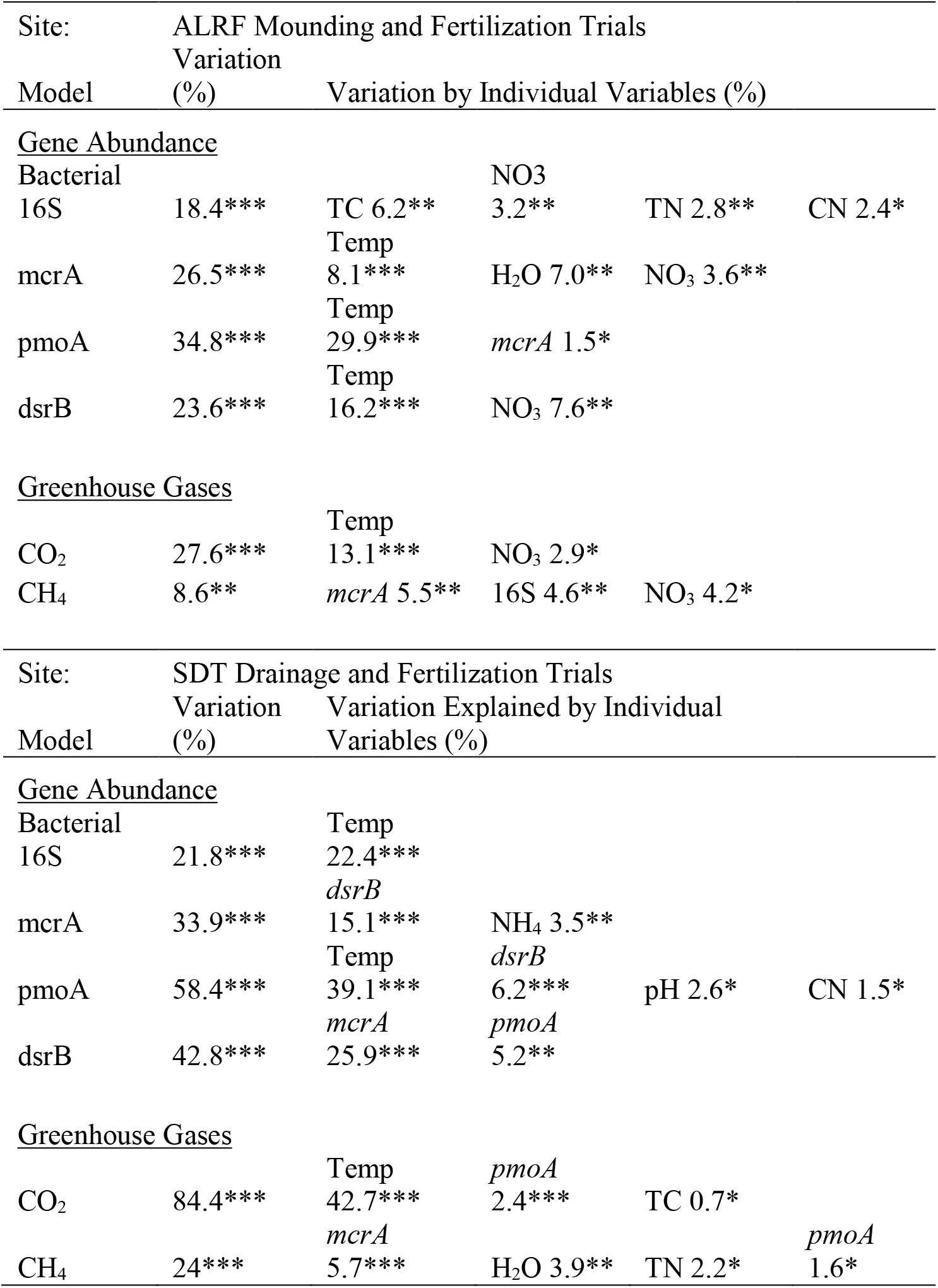
Canonical variance partitioning of gene abundance and greenhouse gas flux rates from ALRF and SDT sites.

|  |  |  |  |  |  |
| --- | --- | --- | --- | --- | --- |
| Site: | ALRF Mounding and Fertilization Trials |  |  |  |  |
|  | Variation |  |  |  |  |
| Model | (%) | Variation by Individual Variables (%) |  |  |  |
| <u>Gene Abundance</u> |  |  |  |  |  |
| Bacterial |  |  | NO3 |  |  |
| 16S | 18.4*** | TC 6.2** | 3.2** | TN 2.8** | CN 2.4* |
|  |  | Temp |  |  |  |
| mcrA | 26.5*** | 8.1*** | H2O 7.0** | NO3 3.6** |  |
|  |  | Temp |  |  |  |
| pmoA | 34.8*** | 29.9*** | mcrA 1.5* |  |  |
|  |  | Temp |  |  |  |
| dsrB | 23.6*** | 16.2*** | NO3 7.6** |  |  |
| <u>Greenhouse Gases</u> |  |  |  |  |  |
|  |  | Temp |  |  |  |
| CO2 | 27.6*** | 13.1*** | NO3 2.9* |  |  |
| CH4 | 8.6** | mcrA 5.5** | 16S 4.6** | NO3 4.2* |  |
| Site: | SDT Drainage and Fertilization Trials |  |  |  |  |
|  | Variation | Variation Explained by Individual |  |  |  |
| Model | (%) | Variables (%) |  |  |  |
| <u>Gene Abundance</u> |  |  |  |  |  |
| Bacterial |  | Temp |  |  |  |
| 16S | 21.8*** | 22.4*** |  |  |  |
|  |  | dsrB |  |  |  |
| mcrA | 33.9*** | 15.1*** | NH4 3.5** |  |  |
|  |  | Temp | dsrB |  |  |
| pmoA | 58.4*** | 39.1*** | 6.2*** | pH 2.6* | CN 1.5* |
|  |  | mcrA | pmoA |  |  |
| dsrB | 42.8*** | 25.9*** | 5.2** |  |  |
| <u>Greenhouse Gases</u> |  |  |  |  |  |
|  |  | Temp | pmoA |  |  |
| CO2 | 84.4*** | 42.7*** | 2.4*** | TC 0.7* |  |
|  |  | mcrA |  |  |  |
| CH4 | 24*** | 5.7*** | H2O 3.9** | TN 2.2* | pmoA 1.6* |

RDA of gene abundances in SDT soil revealed a much stronger correlation between *mcrA* abundance and CH_4_ fluxes at this site (Figure 5b). *McrA* abundance was negatively correlated with SO_4_-S concentrations at this site, yet positively correlated to *dsrB* abundance (Supplementary Table 2). PCNM of SDT sampling locations resulted in 128 PCNM variables between nearest-neighbor sampling locations. Of these, three PCNMs had significant Moran’s I statistics, representing increasingly fine levels of spatial structure. The first PCNM variable at SDT positively correlated with *dsrB, mcrA* and CH_4_ fluxes, these variables exhibiting spatial organization. As with ALRF variation partitioning models, SDT variables were strongly explained by soil temperature (Table 1). 84.4% of CO_2_ flux variation could be explained, with temperature explaining 42.7% on its own. More CH_4_ flux variation was explained at SDT than at ALRF, with *mcrA* abundance again acting as the strongest predictive variable.

## Discussion

CH_4_ flux varies widely in forests associated with different climatic, vegetation and management parameters. In a meta-analysis of CH_4_ flux from UK soils, Levy et al. (2012) report that soil depth, C content, volumetric water content and vegetation cover were the primary site-specific determinants of CH_4_ flux. Site preparation alters many of these influencing factors. For example, Mojeremane et al. (2012) found that drainage of peatland reduced CH_4_ emissions by 57-76%, owing to increased soil temperature and decreased soil water content. Our data generally agree with these observations, and suggest that soil turnover following mounding, and soil water table reduction following drainage, decreases methanogens and dissimilatory SRB.

While it was hypothesized that SO_4_-S may reduce CH_4_ emissions by stimulating SRB to outcompete methanogens (Muyzer and Stams, 2008), there were no correlations between SO_4_-S and CH_4_ (Figure 5). Therefore, there was no evidence from this study that SO_4_-S fertilization at a concentration of 50 kg ha^−1^ can be used to reduce CH_4_ emissions in waterlogged soils. Alternatively, increased CH_4_ efflux is not uncommon in waterlogged forest soils subject to fertilization (Augustin et al., 1998; von Arnold et al., 2005). In a meta-analysis of wetland soil, low N application rates tended to stimulate CH_4_ uptake while larger amounts tended to inhibit uptake by the soil (Aronson and Helliker, 2010). Soil organic C and mineral N availability may also play an important role in regulating CH_4_ flux both in terms of production and oxidation (Krause et al., 2013; Zhuang et al., 2013).

*McrA* and *pmoA* abundance can serve as indicators of the potential for a soil community to emit methane under favourable conditions (Freitag et al., 2010). The abundance of *mcrA* genes unexpectedly responded positively to mineral nitrogen concentrations, suggesting that in nitrogen limited environments methane emissions can be stimulated rather than reduced following fertilization. This result is not without precedent: urea fertilizer can significantly increase CH_4_ emissions (Bodelier, 2011). Basiliko et al. (2009) suggest that in N-limited ecosystems urea-N fertilization can stimulate CH_4_ oxidizing bacteria. Rather than competition between SRB and methanogens for acetate and other substrates, the correlation between *dsrB* and *mcrA* suggests another relationship. Hydrogenotrophic methanogens can syntrophically reduce CO_2_ to CH_4_ using H+ from SRB, allowing both processes to occur in nutrient poor soil (Pester et al., 2012). The recovery of hydrogenotrophic *M. mesophila mcrA* amplicons from soil adjacent to SDT suggests this may be the case, although this hypothesis requires experimental validation. Alternatively, the use of N fertilization levels >100 kg N ha^−1^ yr^−1^ or continuous N deposition can increase CH_4_ fluxes by overwhelming the CH_4_ binding sites in the PMO enzymes in MOB (Basiliko et al., 2009; Jassal et al., 2011; Bodelier, 2011; Gundersen et al., 2012; Zhuang et al., 2013).

Mounding and drainage significantly reduced CH_4_ fluxes after one year. While fertilization significantly increased SO_4_-S concentrations at ALRF, there was no effects on methane rates or methanogen abundance at the conclusion of this study. This work demonstrates that drainage led to a lasting reduction of methanogen populations and CH_4_ emissions, while results were less certain for the mounded site. Therefore, ditch drainage techniques may be a useful method of site preparation to reduce overall GHG emissions from waterlogged, CH_4_-emitting soil. The use of microbial genes abundance can help resolve how the complex changes to the soil community following site preparation can result in alterations to GHG fluxes in waterlogged forest stands.

## Acknowledgements

This work was supported by a Natural Sciences and Engineering Research Council of Canada (NSERC) Collaborative Research and Development (CRD) grant to SJG and CEP (CRDPJ 428948-11) with additional support from Western Forest Products, Inc.; Shell Canada Ltd. provided Thiogro fertilizer. The research was also supported through an NSERC post-graduate award to DL-B. The authors are grateful to Melanie Karjala and the Aleza Lake Research Forest Society for the award of a travel grant which initiated project development. The authors thank Michael Jull for assistance with the installation of mounding trials at Aleza Lake Research Forest, Annette Van Niejenhuis for installation of the Suquash Drainage Trials and Clive Dawson for soil chemical analysis. Jesper Riis Christiansen assisted with gas flux rate analysis. We would also like to thank Charlie Kuan, Angie Li, Eli Rechtschaffen, Leonhard Norz and Ira Sutherland for assistance in the field and laboratory.

---

Figure 5. Redundancy analysis (RDA) of microbial gene abundance (black) by soil variables (blue) and spatial structure (pink) following fertilization. A) Ordination of functional genes constrained by significant soil factors for ALRF samples; b) ordination of SDT functional genes constrained by significant soil factors. Abbreviations: UD, undrained; D, drained; UM, unmounded; M, mounded; F, fertilized (*, p<0.05, **, p<0.01, ***, p<0.001).

## Supplementary Material

**Supplementary Table 1.** Pearson correlation coefficients between measured variables at ALRF.

|  | Microbial genes |  |  |  | GHG emissions |  | Soil characteristics |  |  | Climate |
| --- | --- | --- | --- | --- | --- | --- | --- | --- | --- | --- |
|  | *mcrA* | *pmoA* | *dsrB* | Bact.16S | CO<sub>2</sub> | CH<sub>4</sub> | SO<sub>4</sub>-S | pH | H<sub>2</sub>O | Temp. |
| *mcrA* |  | * |  |  | * | * | *** |  | *** | *** |
| *pmoA* | -0.26 |  |  |  | *** |  |  |  |  |  |
| *dsrB* | 0.09 | -0.05 |  | * |  |  | * |  |  | *** |
| Bact.16S | 0.01 | -0.04 | 0.25 |  | *** |  |  | *** |  | * |
| CO<sub>2</sub> | -0.20 | 0.32 | -0.14 | 0.50 |  | * | * |  |  | *** |
| CH<sub>4</sub> | 0.21 | 0.18 | 0.01 | 0.09 | 0.20 |  |  |  |  |  |
| SO<sub>4</sub>-S | 0.31 | 0.05 | 0.22 | 0.02 | -0.19 | -0.06 |  |  | *** | *** |
| pH | -0.01 | 0.15 | -0.13 | -0.39 | -0.04 | 0.13 | -0.01 |  |  |  |
| H<sub>2</sub>O | 0.41 | 0.00 | 0.14 | 0.19 | 0.06 | 0.00 | 0.37 | -0.16 |  |  |
| Temp. | -0.34 | 0.58 | 0.37 | 0.20 | 0.38 | 0.15 | -0.10 | 0.05 | -0.11 | -0.11 |

(\*, p<0.05, \*\*, p<0.01, \*\*\*, p<0.001)

**Supplementary Table 2.** Pearson correlations between measured variables at SDT.

|  | Microbial genes |  |  |  | GHG emissions |  | Soil characteristics |  |  | Climate |
| --- | --- | --- | --- | --- | --- | --- | --- | --- | --- | --- |
|  | *mcrA* | *pmoA* | *dsrB* | Bact.16S | CO<sub>2</sub> | CH<sub>4</sub> | SO<sub>4</sub>-S | pH | H<sub>2</sub>O | Temp. |
| *mcrA* |  |  | *** |  |  | *** | ** | *** |  |  |
| *pmoA* | 0.03 |  | ** | *** |  | ** |  | ** |  | *** |
| *dsrB* | 0.61 | 0.26 |  |  | ** |  |  | *** |  | ** |
| Bact.16S | 0.09 | 0.38 | 0.01 |  | *** | * |  |  | ** | *** |
| CO<sub>2</sub> | -0.01 | 0.01 | -0.25 | 0.67 |  | * |  |  |  | *** |
| CH<sub>4</sub> | 0.32 | -0.28 | 0.13 | -0.21 | -0.21 |  |  |  | *** |  |
| SO<sub>4</sub>-S | -0.25 | -0.03 | 0.14 | -0.06 | -0.10 | 0.09 |  | -0.43 |  |  |
| pH | 0.48 | 0.27 | 0.32 | 0.13 | -0.03 | 0.02 | -0.43 |  | * |  |
| H<sub>2</sub>O | 0.18 | -0.12 | -0.02 | -0.25 | -0.15 | 0.42 | 0.05 | -0.18 |  |  |
| Temp. | -0.05 | 0.33 | -0.28 | 0.55 | 0.52 | -0.13 | -0.01 | -0.03 | -0.14 |  |

(\*, p<0.05, \*\*, p<0.01, \*\*\*, p<0.001)

